# GENETIC SCREENING OF HALOTHANE GENE ON SELECTED PHILIPPINE NATIVE PIG HERDS

**DOI:** 10.1101/2023.07.07.548076

**Authors:** Sherwin Matias, Maureen Gajeton, Ester Flores

## Abstract

Establishment of nucleus herds (NHs) of Native Pigs (NPs) at various R&D stations in the Philippines is currently being undertaken for food security and genetic conservation. Marker-assisted selection (MAS) is being utilized to identify individuals carrying favorable alleles of genes associated with production traits and screen-out genetic defects (GD) for breeding purposes. Porcine Stress Syndrome (PSS) caused by a mutation in Halothane (HAL) gene is a GD frequently found in commercial breeds that when expressed, causes pale, soft, exudative (PSE) meat. PSE is inferior quality meat undesirable in the market causing economic loss to the swine industry. Thus, this study was conducted to screen HAL gene using mutagenically separated-polymerase chain reaction (MS-PCR) in selected NP herds and assessed its repeatability in local breeds. Results showed that out of 577 screened individuals, 543 (94.11%) were normal (NN), 0 (0%) were homozygous mutant (nn) and 34 (5.89%) were heterozygous carriers (Nn). Therefore, the optimized PSS screening protocol using MSPCR is also applicable to local breed as described in the previous study. As such the availability of genetic test for PSS could be useful in improving the breeding selection and elimination of PSS mutant in the nucleus herd of Philippine Native Pig.

## Introduction

Despite the setbacks and threats imposed by the African Swine Fever (ASF), efforts in establishing nucleus herds of Native Pigs (NPs) are being undertaken at various R&D stations in the Philippines. NPs are known for their valuable traits of being adaptable and resilient to sudden environmental changes. Also, unlike commercial breeds, they only require inexpensive housing facilities, with minimal care, and can be fed with leftovers, vegetable scraps, and plant foliage (Guerero, 2018). Thus, the program encourages the repopulation of this species to promote their market, alleviate food security and strengthen their genetic conservation.

Further, aside from the emergence of diseases, awareness of genetic defects in swine must be heightened too. Porcine Stress Syndrome (PSS) also known as Malignant Hyperthermia (MH) or Halothane Stress Mutation (HSM) is an autosomal recessive disorder that occurred in porcine animals (Iowa State University College of Veterinary Medicine). PSS is caused by a mutation in a single gene called Ryanodine Receptor 1 (RYR1) or Halothane (HAL) Gene. The porcine RYR1 locus was localized on chromosome 6p11-q21 with substitution occurrence at position 1843 (C-T) which corresponds to Arginine-Cysteine shifting at position 615 during translation (Ilie et al., 2014). The RYR1 gene is responsible for channeling the release of calcium ions (Ca^2+^) stored and regulated by the sarcoplasmic reticulum (SR) of the skeletal muscle (Rossi et al., 2008). These Ca^2+^ were being utilized during muscle contraction, relaxation, and energy metabolism (Ilie et al., 2014). Mistranslated amino acids as a consequence of polymorphism cause RYR1 dysfunction which opens and inhibits the closing of the RYR1 channel resulting in the continuous outflow of Ca^2+^ from the SR altering the homeostasis of ions within cells and intracellular (Ilie et al., 2014). Therefore, exposure to stressors and volatile anesthetics like Halothane triggered the onset of PSS.

This HAL gene has two possible alleles, the dominant N allele, and the recessive n allele. These variants are located in a single locus and could transcribe into three possible genotypes which are homozygous normal (NN), heterozygous carrier (Nn), and homozygous mutant/positive (nn) (Manalaysay et al., 2014). The presence of the n allele in hogs is unusually beneficial at the same time a threat to raisers. Advantages of having the n allele include leanness and muscularity in response to hypertrophy (Ilie et al., 2014), this happens usually to heterozygous carriers (Nn). While disadvantageous to hogs having the homozygous mutant genotype (nn) that when exposed to physiological stresses leads to the manifestation of increasing body temperature, muscle rigidity, and metabolic acidosis leading to sudden death and production of pale, soft, exudative (PSE) meat quality postmortem (See et al., 2006).

The first incidence of PSS mutation was recorded in Belgium in Pietrain Breed through intensive selection and subsequent testing (O’Brien 1999). During the period of 1950, the demand for lean and muscled meat was very high despite the poor quality (O’Brien 1999, Stalder and Conatser). A noticeable speed and prevalence of the spread worldwide were attributed to the pyramid structural scheme of the modern swine industry wherein the genetics from the small proportion of the population amplifies to a larger proportion due to the rapid national and international exchange of breeding stocks (O’Brien 1999). Crossbreeding of supposedly purebred breeds of swine also contributed to the widespread of the PSS gene from a single founder breed to numerous breeds.

In the Philippine settings, Manalaysay et al., 2014 optimized the screening protocol of PSS, and it successfully screened PSS incidence in several commercial breeds like Pietrain, Landrace, Large White, Duroc, and Chester white. However, the study was limited to commercial breeds only leaving the Philippine Native pig still unscreened with this genetic condition.

Thus, this study was conducted to screen the prevalence of the HAL gene in Philippine native pig herds. Also, it was conducted to test the repeatability and application of the optimized PSS-screening protocol of Manalaysay et al., 2014 to the local breed. And, determined the incidence of PSS variants by calculating the allelic and genotypic frequencies of each and within samples.

## Materials and methods

### Sample collection

Samples were collected from the different native pig herds inter-island of the Philippines. Sample donors were selected in a completely randomized manner. The housing facility for the animals was mixed type consisting of concrete, metal, and bamboo house pen and some farms practice the free range type of raising. The herds are well-ventilated with the free-flowing air within the area. The ad libitum diets of the animals were mixed with commercial feeds and organic feeding materials like cut and carry grasses and vegetable scraps. The water system in each herd was a combination of deep-well type, barn-fed watering system, and rainwater-fed system.

Samples were collected in the form of blood and tissue (hair follicle and ear fragments). A portable and adjustable animal crate was used to isolate the animal from the herd and to hold them for collection. Using forceps, hair follicles were plucked and put in a labeled zip lock while ear notching was done to collect ear samples. Blood samples, on the other hand, were collected by lifting the head of the animals and performing venipuncture (the needle was punctured at the jugular vein of the animal). The collection kit was composed of a 21 G x 1 1/2” disposable venous blood collection needle, a holder, and a blood collection tube. The tube contains EDTA which acts as an anticoagulant. All samples collected were kept in a cooler with ice packs.

### DNA Extraction

The genomic extraction of tissue (hair and ear) and blood samples was performed using the respective optimized extraction protocol in the Molecular Genetics laboratory. Extraction procedures used were indicated in the kits with minor modifications. For the extraction of tissue samples, Qiagen DNeasy Blood and Tissue Kit was utilized. A tissue (≈ 25 mg for ear fragments and 5-10pcs hair follicles) was cut into pieces and placed in 1.5 microcentrifuge tubes (MCT). Buffer ATL and proteinase K were added and mixed via vortexing. It was incubated at 56°C until it was completely lyzed. Occasional vortexing was performed. Buffer AL was also added and mixed by vortexing. Afterward, it was supplemented with ethanol and subjected to another mixing. To Remove the debris from the tissue and to isolate the genomic DNA, a DNeasy mini spin column was used. Buffer AW1 and AW2 were utilized to wash the sample thoroughly. For the final elution, the spin columns were transferred to a 1.5ml MCT. Buffer AE was used to elute the DNA and it was incubated for 1 minute at room temperature and then stored at 2-8°C.

Conversely, the extraction of blood samples was done using the Promega Wizard Genomic DNA Purification Kit with minor modifications. In a 2ml MCT, 500 ul of homogenized blood samples were transferred. Ammonium chloride (N^4^Cl) was used to wash and pool white blood cells (WBC) from the samples. Cell lysis, nuclei lysis, and protein precipitation were used to isolate the genomic DNA from the WBC. After centrifugation, the supernatant from the mix was transferred to another MCT. Isopropanol was added and mixed by inversion. It was stored overnight and subjected to another centrifugation. For the final washing, 70% ethanol was used and then air-dried inside the laminar flow hood. The DNA samples were eluted using 50 ul of DNA rehydration solution. Tapped the tube to mix the solution then stored at 2-8°C.

### Mutagenically Separated Polymerase Chain Reaction and Gel Documentation

Amplification of the HAL gene was done using mutagenically separated-polymerase chain reaction (MS-PCR). The PCR mix and conditions were based on the optimized protocol of Manalaysay et. al., (2014). The PCR products were verified using 3.5% agarose gel (in 100 ml of TAE add 3.5 g Pronadisa Agarose D1 Low EEO and 3.5 ul Gel Red Biotium Solution) and electrophoresed using Mupid-exU submarine type electrophoresis system. Gel results were viewed using Enduro™ GDS Gel Documentation System. Genotype scoring was performed using standard and manual computation. Sequencing was also done to provide further verification of the gel products and to compare the sequencing result to the findings of Manalaysay et al., 2014 in commercial breeds.

### Statistical Analysis

For the statistical Analysis of this study, a Completely Randomize Design was used and descriptive analysis was done. Hardy-Weinberg Equilibrium Equation was used to determine the Allelic frequency among samples. Genotypic frequency was also computed.

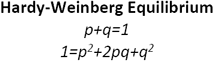

p is the frequency of the dominant allele q is the frequency of recessive allele

p^2^ is the frequency of individuals with the homozygous dominant genotype

2pq is the frequency of individuals with the heterozygous genotype

q^2^ is the frequency of individuals with the homozygous recessive genotype.

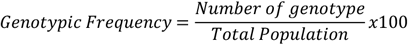

## Results and Discussion

A total of ten (10) sampling sites were coordinated for the sample collection of this study. There were five hundred seventy-seven (577) samples collected and it consist of 285 sows, 158 boars, and 134 unknown sex. Genomic DNA was extracted and genotyped using MSPCR. There were three genotypes determined using the HAL gene, the NN (normal), Nn (carrier), and the (nn) mutant with sizes 114bp, 114 and 134bp, and 134 bp respectively (Figure 1.)

**Figure 1.**
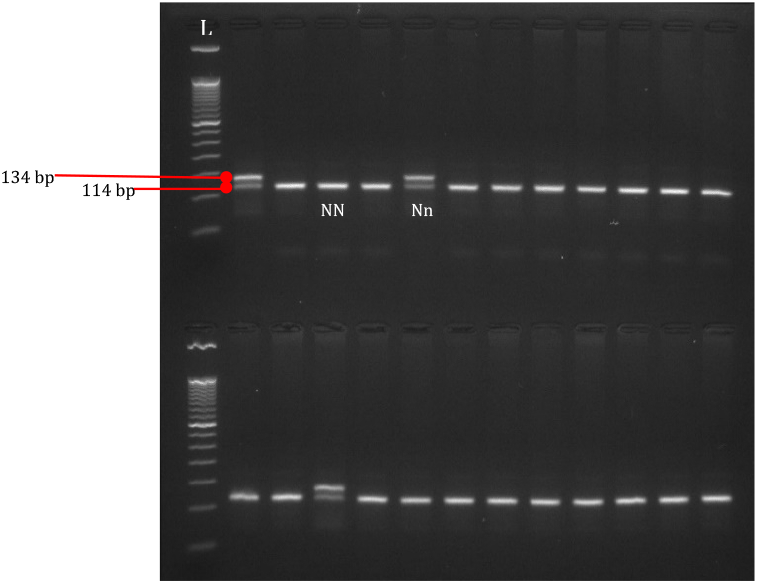
MSPCR gel products of samples having normal (NN) and carrier (Nn) genotypes with 114 bp, 114 and 134 bp sizes resepectively.

Incidence genotypes and genotypic frequency among herds were calculated and tabulated in Table 1. Out of 577 samples, there were 34 carriers (Nn), zero occurrences of mutant (nn), and the rest were normal (NN) with 543 heads. Genotypic and Allelic frequency were also illustrated in Figure 2. Likewise, Figure 3 presented the frequencies of genotypes within respective herds. It was shown that Farms 2, 3,4, 9, and 10 were no incidence of PSS due to zero occurrence of the n allele among the screened samples. While Farm 1, 5,6 7, and 8 screened to have a single copy of the n allele among their samples. Noticeably, Farm 7 has the highest incidence of the PSS variants followed by Farm 5, 6, 1, and 8 (Figure 3, Table 1). As an assumption that all the samples were native pigs, this may imply that the result of this study is the first reported incidence of PSS variants in Philippine Native Pigs. And, the PSS screening protocol as described by Manalaysay et al., 2014, was repeated and applicable to this porcine local species.

**Table 1.** Porcine Stress Syndrome Incidence among selected Native Pigs Herd.

| FARM NO. | N | NORMAL | INCIDENCE % | CARRIER | INCIDENCE % | POSITIVE | INCIDENCE % |
| --- | --- | --- | --- | --- | --- | --- | --- |
| 1 | 105 | 98 | 93.33 | 7 | 6.67 | 0 | 0 |
| 2 | 4 | 4 | 100.00 | 0 | 0.00 | 0 | 0 |
| 3 | 12 | 12 | 100.00 | 0 | 0.00 | 0 | 0 |
| 4 | 51 | 51 | 100.00 | 0 | 0.00 | 0 | 0 |
| 5 | 28 | 25 | 89.29 | 3 | 10.71 | 0 | 0 |
| 6 | 105 | 96 | 91.43 | 9 | 8.57 | 0 | 0 |
| 7 | 96 | 82 | 85.42 | 14 | 14.58 | 0 | 0 |
| 8 | 115 | 114 | 99.13 | 1 | 0.87 | 0 | 0 |
| 9 | 28 | 28 | 100.00 | 0 | 0.00 | 0 | 0 |
| 10 | 33 | 33 | 100.00 | 0 | 0.00 | 0 | 0 |
| **TOTAL** | **577** | **543** | **94.11** | **34** | **5.89** | **0** | **0** |

**Table 2.**
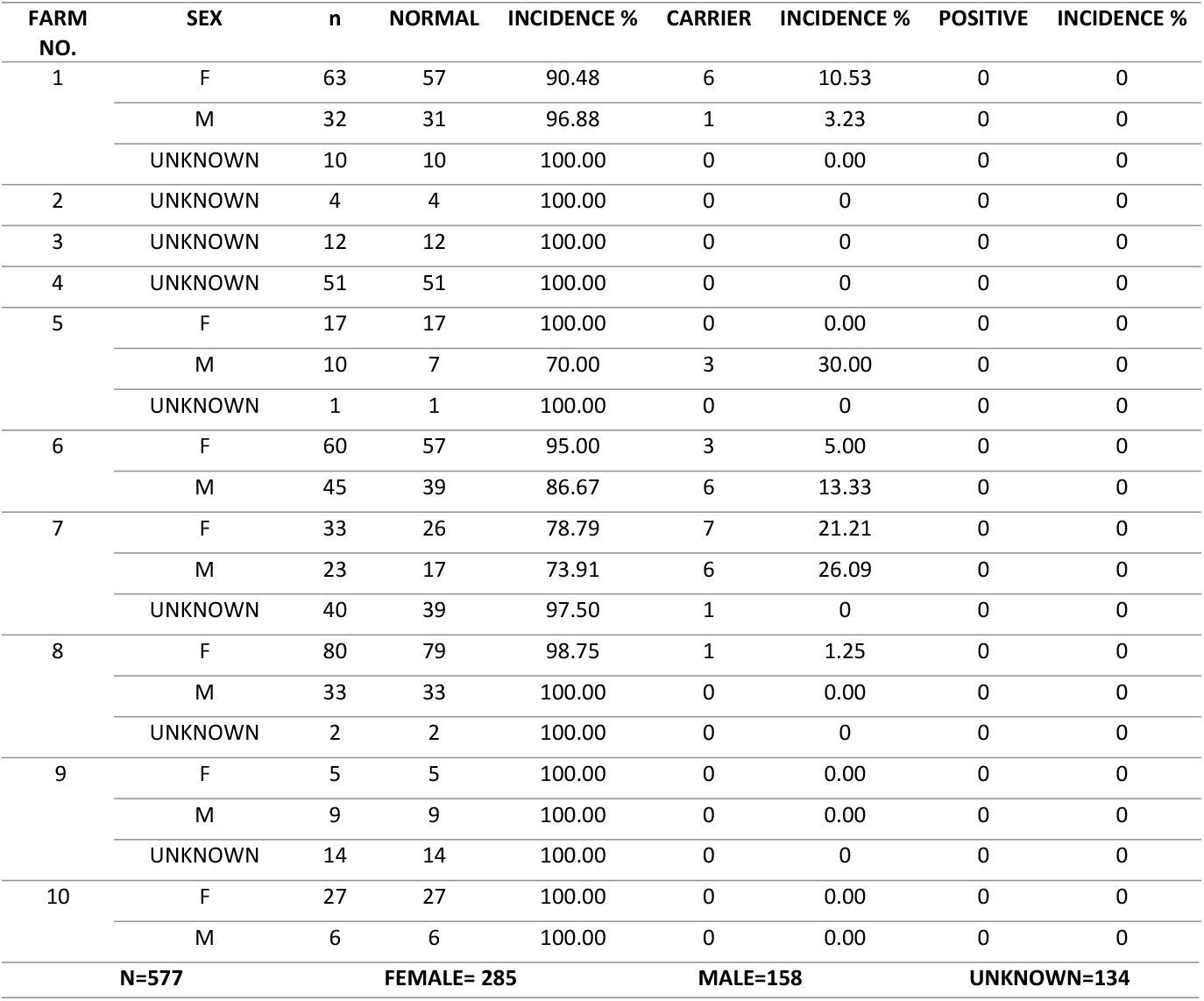
Porcine Stress Syndrome Incidence among selected Native Pigs Herd with regards to sex.

| FARM NO. | SEX | n | NORMAL | INCIDENCE % | CARRIER | INCIDENCE % | POSITIVE | INCIDENCE % |
| --- | --- | --- | --- | --- | --- | --- | --- | --- |
| 1 | F | 63 | 57 | 90.48 | 6 | 10.53 | 0 | 0 |
|  | M | 32 | 31 | 96.88 | 1 | 3.23 | 0 | 0 |
|  | UNKNOWN | 10 | 10 | 100.00 | 0 | 0.00 | 0 | 0 |
| 2 | UNKNOWN | 4 | 4 | 100.00 | 0 | 0 | 0 | 0 |
| 3 | UNKNOWN | 12 | 12 | 100.00 | 0 | 0 | 0 | 0 |
| 4 | UNKNOWN | 51 | 51 | 100.00 | 0 | 0 | 0 | 0 |
| 5 | F | 17 | 17 | 100.00 | 0 | 0.00 | 0 | 0 |
|  | M | 10 | 7 | 70.00 | 3 | 30.00 | 0 | 0 |
|  | UNKNOWN | 1 | 1 | 100.00 | 0 | 0 | 0 | 0 |
| 6 | F | 60 | 57 | 95.00 | 3 | 5.00 | 0 | 0 |
|  | M | 45 | 39 | 86.67 | 6 | 13.33 | 0 | 0 |
| 7 | F | 33 | 26 | 78.79 | 7 | 21.21 | 0 | 0 |
|  | M | 23 | 17 | 73.91 | 6 | 26.09 | 0 | 0 |
|  | UNKNOWN | 40 | 39 | 97.50 | 1 | 0 | 0 | 0 |
| 8 | F | 80 | 79 | 98.75 | 1 | 1.25 | 0 | 0 |
|  | M | 33 | 33 | 100.00 | 0 | 0.00 | 0 | 0 |
|  | UNKNOWN | 2 | 2 | 100.00 | 0 | 0 | 0 | 0 |
| 9 | F | 5 | 5 | 100.00 | 0 | 0.00 | 0 | 0 |
|  | M | 9 | 9 | 100.00 | 0 | 0.00 | 0 | 0 |
|  | UNKNOWN | 14 | 14 | 100.00 | 0 | 0 | 0 | 0 |
| 10 | F | 27 | 27 | 100.00 | 0 | 0.00 | 0 | 0 |
|  | M | 6 | 6 | 100.00 | 0 | 0.00 | 0 | 0 |
| **N=577** |  | **FEMALE= 285** |  |  | **MALE=158** |  | **UNKNOWN=134** |  |

**Figure 2.**
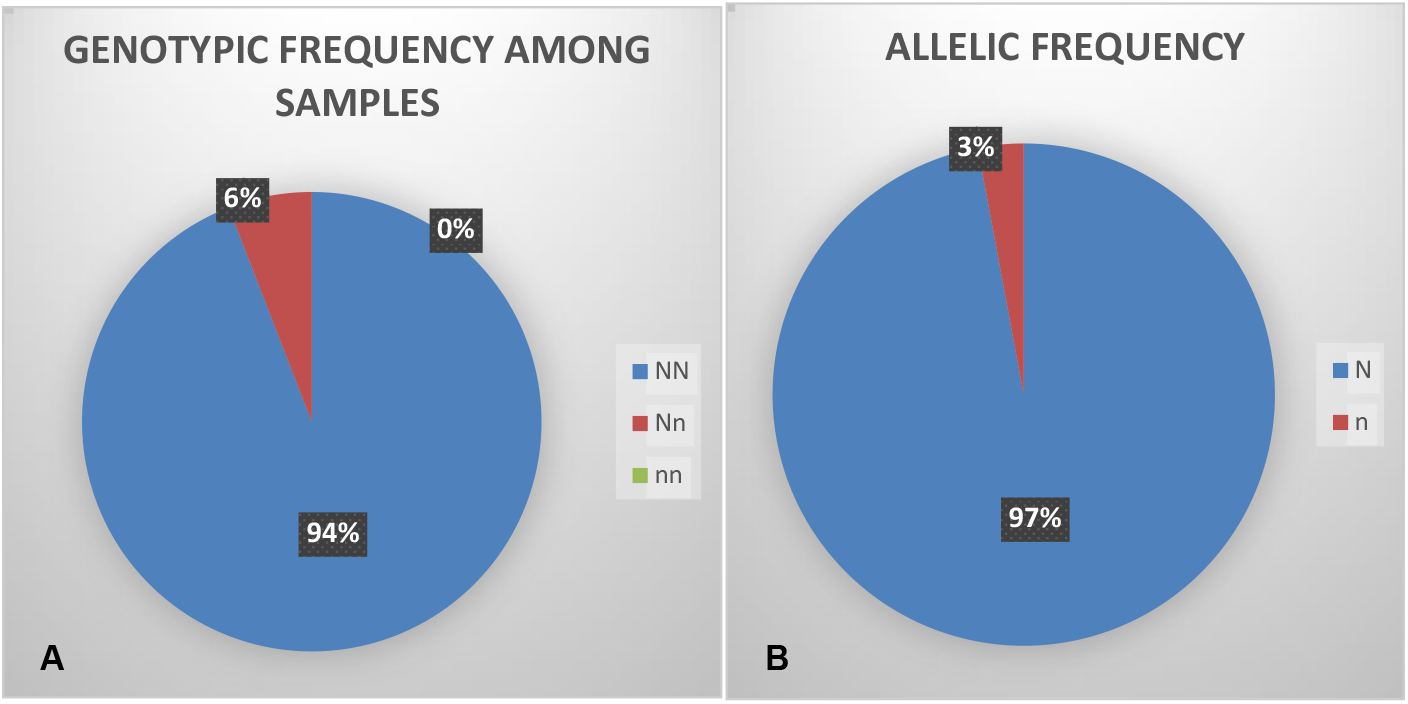
A.) Genotypic frequency among samples B.) Allelic Frequency.

**Figure 3.**
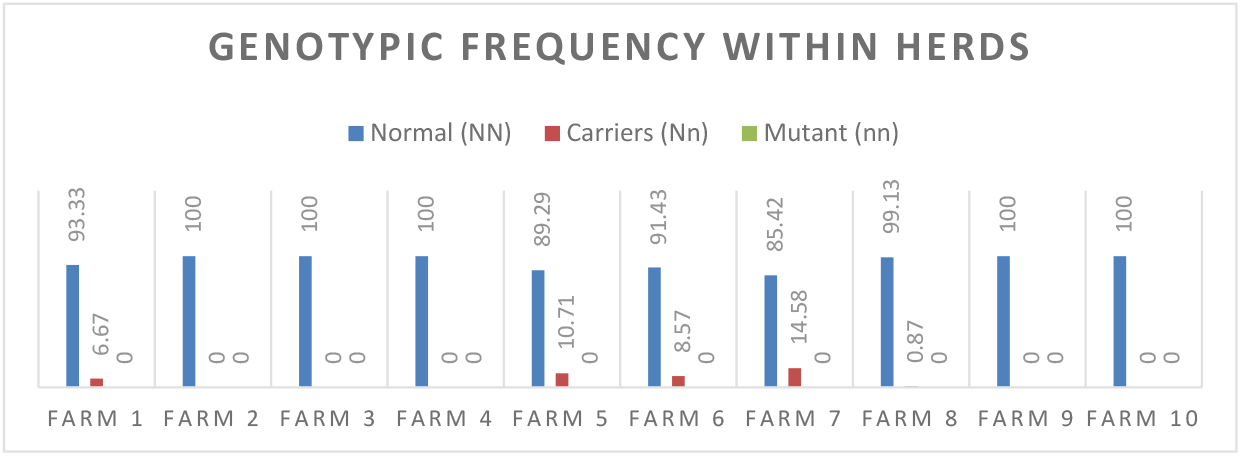
Graphical representation of genotypic frequency within herds.

In general, prompting genetic defects carrier into the population is sometimes subjected to culling because it has a perceivable threat to the entirety of the population. However, in the case of the PSS mutation, it has a binary outturn that gives options to raisers on whether to utilize or eliminate the variant. It is given that the presence of the single copy of the n allele is associated with meat quality such as lean, heavy muscling, and increased growth efficiency (O’Brien 1999) which are advantageous traits to raisers. On the contrary, mutant/PSS-positive (nn) could lead to deleterious effects such as stress-susceptibility producing poor quality meat (Stalder and Conatser 1999; O’Brien 1999) that may have caused losses to them. Providentially, this negative outturn is highly variable and could be modified by environmental and management factors (O’Brien 1999, Stalder and Conatser). Also, the molecular level of analysis in the detection and selection of such is now very advance, accurate, and with a lesser turnaround of results. Thus, elimination of such genotype can be done as early as possible and could give chance in retaining the desirable traits alone.

## Conclusion

The study, therefore, concluded that there was screened single copy of the n allele in Philippine native pigs. Thus, there is a possibility of having or could have had the mutant genotype. Also, the optimized PSS screening protocol of Manalaysay et al., 2014 has been tested and repeated for the result’s accuracy and reliability to local breed.

## Acknowledgments

This study was conducted in collaboration with the Philippine Carabao Center and various R&D Stations and SUC’s in the Philippines. This program is funded by the DOST-PCAARRD.

